# Evidence of physiological assortment and movement dynamics among social groups of a coral reef fish

**DOI:** 10.1101/2025.07.09.663861

**Authors:** Lauren E. Nadler, Mark I. McCormick, Amy Cox, Kathryn Grazioso, Shaun S. Killen

## Abstract

The trade-offs of group living are modulated by the phenotypes of individual members of a social group, particularly in dynamic and diverse habitats like coral reefs. Little is known about the patterns of physiological traits among fishes within social groups and the mechanisms that promote these patterns, which could elucidate the drivers of group composition and their downstream ecological and evolutionary impacts. Here, in the gregarious damselfish species *Chromic viridis*, we examined inter-group differences in whole-animal physiological traits and the tendency for fish to move either within sites (i.e., sections of continuous reef) or among habitats (i.e., reefs separated by sandy substratum) to a new social group. Using oxygen uptake as a proxy for aerobic metabolic rate, we found significant differences in maximum metabolic rate (MMR) and aerobic scope (AS) among schools from different habitats, with these traits higher in habitats with faster water flow rates. However, we found no differences in any metabolic traits (standard metabolic rate, SMR, MMR, AS) between groups from the same site.

These trends could stem from a range of mechanisms, as mark-recapture studies of this species indicated a willingness to migrate to a new social group in over 30% of recollected fish. However, there were no effects of either body size or perceived habitat risk on the distance moved or movement type (i.e., over coral or sand). Our results indicate that, in social species, a combination of mechanisms may influence phenotypic differences among groups over different spatial scales.

## Introduction

In social animal groups, group phenotypic composition greatly influences how individuals experience the trade-offs of group living (Jolles et al. 2020). Although living in a group increases foraging success, predator avoidance and efficiency of movement, group living animals also experience greater competition for resources, agonistic encounters and exposure to novel infectious pathogens (Krause and Ruxton 2002; Ward and Webster 2016). To maximize the benefits and minimize the costs of sociality, individuals may choose to socialize with conspecifics that are similar to themselves, with previous studies illustrating similarities in traits like body size and behaviour among individuals in social animal groups (Rodgers et al. 2011; Cattelan and Griggio 2018; Kelley and Evans 2018; Darden et al. 2020). However, although these traits are all correlated with numerous whole-animal physiological traits, including metabolism, immunity, growth and endocrine status (Stamps 2007; Koolhaas 2008; Killen et al. 2016b; Lopes 2017), the role of individual physiology in group phenotypic composition in natural habitats has thus far remained largely unexplored.

Similarity in appearance, body size or behaviour could act as proxies for similarity in physiological traits or performance capacity, particularly those related to energy metabolism (Metcalfe et al. 2016). Resting metabolic rate (standard metabolic rate, SMR, for animals at a particular temperature; basal metabolic rate in thermoneutral endotherms; Metcalfe et al. 2023), for example, has been linked to sociability and risk-taking behaviour (McKenzie et al. 2015; Killen et al. 2016a). Variation in maximum metabolic rate (MMR; upper constraint on an individual’s oxygen-consuming physiological activities) and aerobic scope (AS; capacity to deliver oxygen to support aerobically-fuelled activities beyond basic maintenance) are related to differences in the capacity for aerobic locomotion among individuals and tolerance for environmental stressors (Claireaux and Lefrancois 2007; Killen et al. 2013). Therefore, actively choosing (i.e., through active assortment) to socialize with physiologically-similar conspecifics may be beneficial, as group-mates would exhibit similar nutritional and habitat requirements, tolerance for environmental conditions, and locomotor abilities (Killen et al. 2017b). Thus, homogeneity in physiological traits could provide selective advantages to group-living animals (Jolles et al. 2020), as it would maximize the chance of living in a suitable habitat and aid in minimizing the oddity effect, in which predators preferentially target individuals that stand out from the rest of the group (Landeau and Terborgh 1986; Ruxton et al. 2007). Despite the theoretical background to support selective advantages of physiological assortment among animal groups in the wild (Killen et al. 2017b), these ideas have never been empirically investigated.

Spatial variability in morphological, behavioural, and physiological phenotypes is likely shaped by a range of non-exclusive mechanisms. First, phenotypic plasticity could shape the traits expressed in individuals, as a single genotype can produce multiple alternative phenotypes in response to environmental conditions (Chevin and Hoffmann 2017). In addition, when selective pressures on particular phenotypes vary in different habitats, we would expect to see selective mortality, in which specific phenotypes increase an animal’s vulnerability to predation (Higham et al. 2015). Connections between physiology and habitat preferences (e.g., links between metabolic rate, growth, and thermal preferences, Killen 2014) could also lead to spatial and temporal overlap in specific phenotypes and promote homogeneity within social groups through passive assortment mechanisms. As such, these mechanisms may lead to heterogeneity in the distribution of phenotypes both within and among social groups from different habitats (Killen et al. 2017b). In fishes, water flow has been linked to differences in behavioural and physiological phenotypes relevant for effective social behaviour. Under higher water flow conditions, fishes exhibit faster escape responses (Nadler et al. 2018), greater swimming capacity (Fulton et al. 2013; Binning et al. 2014) and higher aerobic performance (Binning et al. 2015). Therefore, water flow could be a contributing factor driving the distribution of physiological traits or performance capacity within and among social groups of fishes (referred to as “schools” from here on), but this has yet to be studied in natural systems.

There are also events that cause group composition to change as individuals are exchanged among social groups (Aureli et al. 2008). While species are known to vary in their propensity for fission (i.e., splitting) and fusion (i.e., merging) events, several ecological pressures and social traits will modify the trade-offs of fidelity to a single school or migration to a new school. Food availability is likely to alter group fidelity, with individuals coping with sparse or inconsistent food resources through flexible social associations (Zheng and Fu 2021). Alternatively, risk from consumers like predators and parasites may de-incentivize migration among schools, due to the risk associated with migration. Kelley et al. (2011) found that shoaling guppies (*Poecilia reticulata*) maintain stronger and more consistent social associations when experiencing high predation risk than habitats with a lower risk profile. Factors associated within the shoal itself can also modify the trade-offs of shoal fidelity versus migration, such as group size, density, familiarity, and personality of group members (Lee-Jenkins and Godin 2010; Cote et al. 2012; Bierbach et al. 2020; Durrer et al. 2020). For some of the physiological assortment mechanisms that are outlined above to occur (e.g., active choice), it would require the species willingness and ability to migrate to a group or habitat with preferred characteristics. However, whether that migration is occurring in dynamic and inherently risky habitats like coral reefs remains poorly understood.

Coral reefs are dynamic and vibrant habitats, which drives diverse forms of social grouping in coral reef fishes, including both roving and site-attached species as well as those that live in the water column above the reef or are reliant on the reef structure as an integral component of their defensive strategy against predators (Welsh and Bellwood 2012; Engel et al. 2024). This diversity is driven in part by the challenging environmental conditions found on coral reefs, including rapid water flow rates and unpredictable water turbulence (Clarke et al. 2009; Johansen 2014). Further, predation risk on coral reefs is extensive, due to the diversity of hunting modes and density of predators, meaning prey need to effectively use a range of antipredator strategies to survive (McCormick et al. 2018; Palacios et al. 2018). Many species match their lifestyle to the unique pressures of a particular life stage, partaking in cooperative social grouping as vulnerable juveniles. For example, the parrotfish *Chlorurus sordidus* school in the juvenile phase when they are undergoing rapid colour changing, preferentially schooling with other individuals with which its colouration matches and shifting social associations as needed as colouration changes to mitigate predation risk (Crook 1999b,a). Given the unique challenges that social fishes face on coral reefs, the pressures for optimal social group composition may be high, creating an ideal natural laboratory to investigate physiological assortment and movement dynamics in social fishes.

The present study examined this issue in wild schools of a gregarious coral reef damselfish, the blue-green chromis (*Chromis viridis*). Our objectives were to: 1) to establish whether there were differences in metabolic traits (SMR, MMR and AS) among wild schools within and among habitats; and 2) to determine the frequency and type of movement of individuals among social groups in this species to understand the potential mechanisms driving differences in metabolic rate among social groups. We hypothesized that characteristics like MMR and AS, which are subject to plasticity through a training effect and selection due to their role in predator avoidance (Davison, 1997; Killen et al., 2015), would vary spatially with physical environmental conditions (i.e., among habitats). Conversely, SMR influences energetic and nutritional requirements (Killen et al., 2016b), and therefore, may promote assortment of individuals on a local scale (i.e., within sites) due to differences in resource requirements. The propensity to migrate to new social groups may increase in individuals with larger relative body size and occur with greater frequency when individuals perceive a greater level of threat in their habitat.

## Materials and methods

### Study 1: Physiological differences among social groups

#### Study site and specimen collection

This study was conducted at the Lizard Island Research Station (LIRS) in the northern Great Barrier Reef, Australia (14°40′S, 145°28′E), from November to December 2013. Wild schools of the damselfish *C. viridis* (a live-coral associated schooling species with high site fidelity; n=11 groups total) were collected from shallow coral reef habitats (<4 m, Figure 1) surrounding LIRS using a barrier net, hand nets and a dilute anaesthetic solution composed of clove oil, ethanol and seawater (Munday and Wilson, 1997). This combination of collection materials and techniques allowed most of the individuals in each school to be collected. Each school contained approximately 50 to 100 fish.

**Figure 1.**
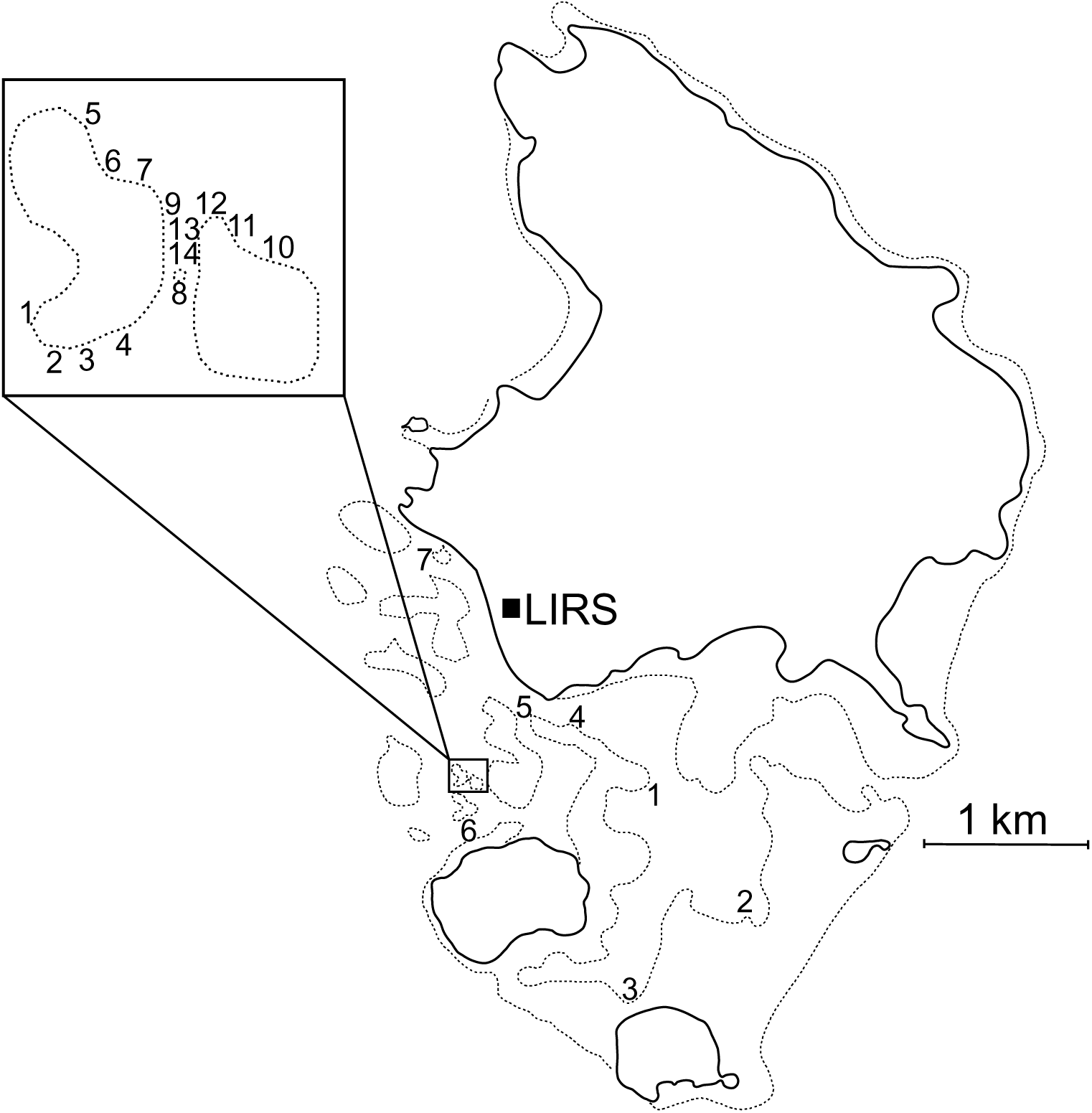
Map of the study site, Lizard Island (LIRS = Lizard Island Research Station). The numbers in the large map show the seven sites around Lizard Island from which individuals from wild *Chromis viridis* schools were collected for experiment 1. The inset map indicates the locations of *C. viridis* schools that were tagged with visible implant elastomer for experiment 2. Solid lines denote land mass, while dotted lines indicate the location of shallow coral reef habitats around Lizard Island.

The number of schools that were collected within a particular shallow reef habitat was dictated by the availability of suitable schools that were >50m apart (in an effort to ensure that they were distinct from each other). At three habitats, one school was collected; at four habitats, two separate groups were collected. Schools within a habitat were collected on the same day. Habitats were separated by 400 – 3000 m and occurred on non-continuous sections of reef (i.e., fish would need to cross a section of sandy substrate to move between habitats but not between two schools at the same habitat). Each of the habitats (i.e., sections of continuous reef) where multiple schools were collected exhibited consistent bathymetry between the locations where schools were collected.

Following collection, schools were returned to the wet laboratory at LIRS. Within a single school, *C. viridis* can exhibit high variability in body size (∼1-5 cm) and all the sampled schools exhibited comparable body size distributions (as determined through qualitative field observations of groups on SCUBA). For testing, given the strong influence of body mass on metabolic rate (Clarke and Johnston 1999), eight individuals from an intermediate size class that was represented in all of the wild schools (3.0 – 3.7 cm standard length) were chosen randomly for measurement of metabolic traits, allowing us to look at relative differences in mean metabolic rate in a comparable subset of each school. In total, 88 fish were measured from the 11 schools collected, which accounted for approximately 10% of all fish in these wild schools. Prior to measurement of metabolic traits (which took place three to five days after collection from the reef), schools were housed in 15L flow-through aquaria (37 x 22 x 20 cm, L x W x H) under natural light conditions and ambient temperature. Fish were fed to satiation twice daily with INVE Aquaculture pellets and newly hatched *Artemia* sp.

#### Respirometry

Metabolic rate of individual fish was estimated using intermittent-flow respirometry, a technique in which oxygen uptake is measured as a proxy for aerobic metabolism (Svendsen et al. 2016). The 53 essential criteria for reporting of methods in intermittent-flow respirometry are included in the supplementary material (Table S1; Killen et al. 2021b). Prior to testing, food was withheld for 24 hours to ensure that fish were in a post-absorptive state (Niimi and Beamish 1974). Maximum metabolic rate (MMR) was then measured by manually stimulating fish to swim in a 40 cm-diameter circular container filled to a depth of 20 cm, until the fish no longer responded to chasing with burst swimming (mean time = 90.07 s ± 3.32 s.e.m; Killen et al. 2017a). Following this chase protocol, fish were air-exposed for 30s to further ensure that they had depleted all endogenous oxygen stores. They were then transferred to individual cylindrical glass respirometers with acrylic end-caps (total volume of chamber plus associated tubing = 75 ml). This established method measures the maximum rate of oxygen uptake during the aerobic recovery period following exhaustive anaerobic exercise, which a broad comparative meta-analysis indicated is comparable to MMR induced by prolonged swimming in most fishes (Killen et al. 2017a).

Respirometers were immersed in a water bath, which was supplied with water from a header tank in which temperature was controlled to 28.74°C (± 0.14 °C, mean ± s.e.m.). Fish were left undisturbed in respirometers overnight and in complete darkness for the next 14 h. Opaque dividers were placed between adjacent respirometers to prevent visual stimulation of activity between fish. Water flow from the temperature-controlled header tank through the respirometers was driven by an external pump in an adjacent header tank set to alternatingly turn on (2 min) and off (7 min) throughout the measurement period. This timing allowed decreases in water oxygen content to be measured every 2 s for 7 min while the respirometer was in the closed state, and then the respirometer was flushed with aerated water for 2 min. An exception was the first closed phased after the chasing protocol to elicit MMR, during which the chamber was closed for at least 10 min to ensure that the immediate post-exercise phase was measured for oxygen uptake. Water mixing within each respirometer was achieved with a peristaltic pump that moved water through the chamber and around an external circuit of gas-impermeable tubing. Also located within the circuit for each respirometer was a flow-through cell, which housed an oxygen-sensing optode attached to an oxygen sensor (Firesting 4-Channel oxygen meters; Pyroscience, Germany) and computer.

Slopes were calculated from the plots of oxygen concentration versus time using linear least squares regression, and then converted to determine the rate of oxygen uptake (mg O_2_ h^-1^). All R^2^ values were greater than 0.95. The first minute of each closed phase was excluded from analyses to ensure adequate mixing within the chamber and external circuit. MMR was calculated by analysing the first slope after exhaustive exercise, and then taking the highest 3-min time interval recorded during this time as MMR. SMR was taken as the lowest 10^th^ percentile of measurements excluding the first 2 h after exercise. Rates of oxygen uptake were elevated during this time (Norin and Clark 2016) but reached a stable level within 2 hours after exercise. AS was calculated as the difference between MMR and SMR. To correct for background bacterial oxygen consumption, oxygen uptake in all chambers was measured while empty both before and after each trial, and all measures of oxygen uptake were corrected assuming a linear increase in bacterial oxygen consumption with time (Rodgers et al. 2016). Each morning, all respirometers, flow-through cells, and tubing were thoroughly cleaned with soap, bleach and hot water, which aided in maintaining background respiration at a mean of 17.03% of SMR (± 1.20% s.e.m).

#### Environmental Variables

Water temperature, depth, and mean water flow rate were measured at each fish collection habitat. As neither water temperature (as measured using a thermometer) nor depth (as measured using a dive computer) exhibited variability among habitats (i.e., within 1°C during all measurements and the depth range was only 1.8 to 4.0 m), neither temperature nor depth were included in any further analyses. Water flow rate was measured on five separate days (under varying wind and weather conditions) throughout December 2013. These measurements allowed us to look at relative differences in conditions among habitats. Measurements were always recorded during daylight hours and at high tide (± 1 hour). Flow rate at each location was determined using a precision vane-wheel flow meter (Hontzsch Gmbh, Waiblingen, Germany) that was placed into the flow approximately 1.25 m below the surface. As *C. viridis* forages on plankton in the water column above the reef during the day (Chase et al. 2020), this depth would be a realistic indicator of the flow conditions experienced by these schools during processes that require swimming (particularly foraging and escaping predators). Measures of flow speed (cm s^-1^) were recorded at 1 Hz for 180s, which was used to calculate each habitat’s mean flow rate during each measurement. An overall mean flow rate was then calculated for each habitat using data for all five days (with s.e.m. within habitats varying from 1.2 to 8.2 cm/s). All measurements were conducted at wind speeds <15 knots (as is typical on most days in the area in the spring/summer season), as previous work at Lizard Island indicates that water flow at habitats comparable to ours (shallow, sheltered reef sites) are consistent through time under these conditions (Johansen 2014), allowing us to look at relative differences among habitats.

### Study 2: Mark-recapture of social groups

This study was conducted at the LIRS from November to December 2015. On two shallow coral reef habitats adjacent to the LIRS (Figure 1, Little Vickis, No Name reefs, sand between Little Vickis and No Name reefs), individuals from fourteen schools of *C. viridis* (all schools contained > 100 individuals) were collected using hand nets and barrier nets (5 mm mesh size). These two reefs were isolated from each other by a stretch of sandy substrate, measuring a minimum of 13 m. As *C. viridis* is an obligate branching coral dweller that sleeps amongst coral branches at night (Chase et al. 2020), the corals on which these schools were found were assigned a number and tagged with fluorescent flagging tape so that they were identifiable on subsequent visits. From each marked coral, 20 individuals were collected and tagged with a unique colour combination of visible implant elastomer (VIE) on SCUBA. Where possible, these 20 individuals were an even mix of small (< 4.0 cm SL) and large (> 4.0 cm SL) body sizes. Two of the schools collected were predominantly one of these size classes (only 5 large individuals captured from school 7 and only 4 small individuals captured from school

10). In these cases, additional individuals of the predominant size class were tagged, so that all schools contained 20 tagged individuals. After 9 – 11 days following the initial collection, 7 of the 14 schools were recollected and the tag and size class of all marked fish were recorded. Then, 20-22 days after the first collection, all schools were recollected, and size and tag characteristics were recorded for all marked fish.

Recollection was staggered in this way, so that the effect of collection and hence perceived risk of the environment on fish migration could be determined.

#### Statistical analyses

All analyses and figures were completed with R v.4.4.1 (R Development Team 2024), using the packages “glmmTMB” (Brooks et al. 2017), “emmeans” (Lenth 2024), “DHARMa” (Hartig 2022), “car” (Fox and Weisberg 2019), “performance” (Lüdecke et al. 2021) and “ggplot2” (Wickham 2016). The level of significance for all tests was α = 0.05. Normality, linearity and homogeneity of residuals were verified by inspection of residual- fit plots and qq-norm plots using the DHARMa package. To determine the best-fit model, we completed model comparison using Akaike’s Information Criterion corrected for small sample size (AICc; Burnham and Anderson 2002).

For study 1, we first examined the effect of water flow rate on metabolic traits. Flow measurements indicated a strongly bimodal distribution of flow values among sites (Table 1), and so water flow rate was designated as a categorical variable, with flow > 20 cm s^-1^ on average designated as ‘high flow’ and all other flows (all < 11 cm s^-1^) designated as ‘low flow’. The effect of water flow on the metabolic traits were examined using three generalized linear mixed-effects models (GLME), with raw SMR, MMR, or AS (all mg O_2_ h^-1^) as the dependent variable, body mass (g) as a co-variate, water flow rate and its interaction with body mass as fixed effects, and school as a random effect. We then examined differences between individuals within habitats where two schools were collected. For these GLMEs, SMR, MMR, or AS (all mg O_2_ h^-1^) was the dependent variable, body mass (g) was a covariate, school and its interaction with body mass were fixed effects, and site was a random effect. A Tukey’s multiple comparisons post-hoc test between schools within sites was used for any model with a significant school effect, to determine if schools within sites showed significant differences in any metabolic traits.

**Table 1.** Characteristics of the sites sampled in study 1, with the mean body mass for sampled fish within schools at each site. The column ‘map label’ refers to the corresponding label number in Figure 1. The columns “Body mass” and “Standard length” refers to the mean mass and length of the fish from each school that were used in the study (n=8 fish per school). Error = s.e.m.

| Site | School | Map label | Coordinates | Flow (cm s<sup>-1</sup>) | Temperature (°C) | Depth (m) | Body mass (g) | Standard length (cm) |
| --- | --- | --- | --- | --- | --- | --- | --- | --- |
| Channel | 1 | 1 | S 14° 41.052', E 145° 26.938' | 21.8±8.3 | 28.6±0.2 | 1.8 | 1.21±0.07 | 3.23±0.22 |
| Channel | 2 | 1 | S 14° 41.052', E 145° 26.938' | 21.8±8.3 | 28.6±0.2 | 2 | 1.60±0.09 | 3.34±0.16 |
| Loomis | 3 | 2 | S 14° 41.125', E 145° 27.020' | 21.8±8.4 | 28.5±0.3 | 2.3 | 1.67±0.06 | 3.24±0.19 |
| Loomis | 4 | 2 | S 14° 41.125', E 145° 27.020' | 21.8±8.4 | 28.5±0.3 | 2.3 | 1.78±0.07 | 3.60±0.18 |
| Lagoon | 5 | 3 | S 14° 41.363', E 145° 27.363' | 8.4±1.57 | 28.1±0.3 | 2.5 | 1.75±0.09 | 3.47±0.11 |
| Lagoon | 6 | 3 | S 14° 41.363', E 145° 27.363' | 8.4±1.57 | 28.1±0.3 | 2.5 | 1.85±0.20 | 3.48±0.20 |
| Bird Islet | 7 | 4 | S 14° 41.885', E 145° 27.651' | 10.0±1.8 | 28.1±0.2 | 4 | 1.82±0.09 | 3.46±0.12 |
| Bird Islet | 8 | 4 | S 14° 41.885', E 145° 27.651' | 10.0±1.8 | 28.1±0.2 | 3.6 | 2.05±0.09 | 3.63±0.09 |
| Horseshoe | 9 | 5 | S 14° 41.387', E 145° 26.689' | 10.0±2.2 | 28.5±0.2 | 2.2 | 1.576±0.076 | 3.59±0.125 |
| Station | 10 | 6 | S 14° 40.845', E 145° 26.746' | 8.2±1.2 | 28.2±0.2 | 2.0 | 1.677±0.089 | 3.52±0.168 |
| South Island | 11 | 7 | S 14° 42.022', E 145° 27.173' | 9.0±1.3 | 28.1±0.2 | 3.4 | 1.955±0.108 | 3.37±0.082 |

For study 2, we first examined factors influencing the tendency to move (yes, no) using a GLME with a binomial distribution, with body size class (small, large), recapture frequency (of the school in which the fish was recaptured; once, twice), and their interaction as fixed effects, with school included as a random effect. Then, in the subset of the data for fish that migrated, we examined the distance moved (m) and movement type (coral, sand; modelled with a binomial distribution). For these GLMEs, we included body size class, recapture frequency (of the original school), and their interaction as fixed effects, with school included as a random effect. We also conducted a network analysis using the mark-recapture data using the igraph R package, for network construction and metrics calculations (Csardi and Nepusz 2005; Csárdi et al. 2024).

Each colony was represented as a node, with directed edges between nodes indicating movement of individual fish from one colony to another. The weight of each edge was defined by the number of fish moving in that direction, representing movement intensity among colonies. To ensure that only inter-colony movements were analysed, we excluded instances where fish were recaptured within the same colony. To quantify colony connectivity and movement dynamics, we calculated the following network metrics for each colony: 1) in-degree: representing the total number of fish arriving at a colony from other colonies. Higher in-degree values suggest colonies that are more attractive or central for movement among colonies. 2) out-degree: representing the total number of fish leaving a colony to other colonies. This metric indicates colonies with higher dispersal rates. 3) total degree: the sum of in-degree and out-degree, an overall measure of connectivity for each colony as either a source or destination. 4) betweenness: quantifies the role of a colony as an intermediary or "bridge" in the network. Colonies with higher betweenness may connect different parts of the network. 5) eigenvector centrality: summarises direct connections of a colony and the centrality of its neighbours, allowing colonies with connections to other well-connected colonies to have higher values. We investigated how node metrics (degree, eigenvector centrality) change with recapture frequency and site (Little Vickis, No Name, sand between Little Vickis and No Name reefs) using a generalized linear model (GLM).

## Results

### Study 1: Physiological differences among social groups

The results indicated significant variation in some metabolic traits among habitats, linked to each habitat’s respective flow rate. Characteristics of the study sites and mean body mass for fish within schools at each site are summarized in Table 1. The best-fit models for MMR and AS included water flow rate, body mass, and their interaction, but did not include a site random effect. Both MMR (Figure 2b) and AS (Figure 2c) were higher for fish in schools exposed to high flow habitats (MMR: χ^2^=8.06, p = 0.005, adjusted R^2^=0.401; AS: χ^2^=7.46, p = 0.006, adjusted R^2^=0.311). This effect was enhanced by larger body mass, as MMR and AS also exhibited a significant water flow rate*body mass interaction (MMR: χ^2^= 3.97, p = 0.046; AS: χ^2^= 4.17, p = 0.041).

**Figure 2.**
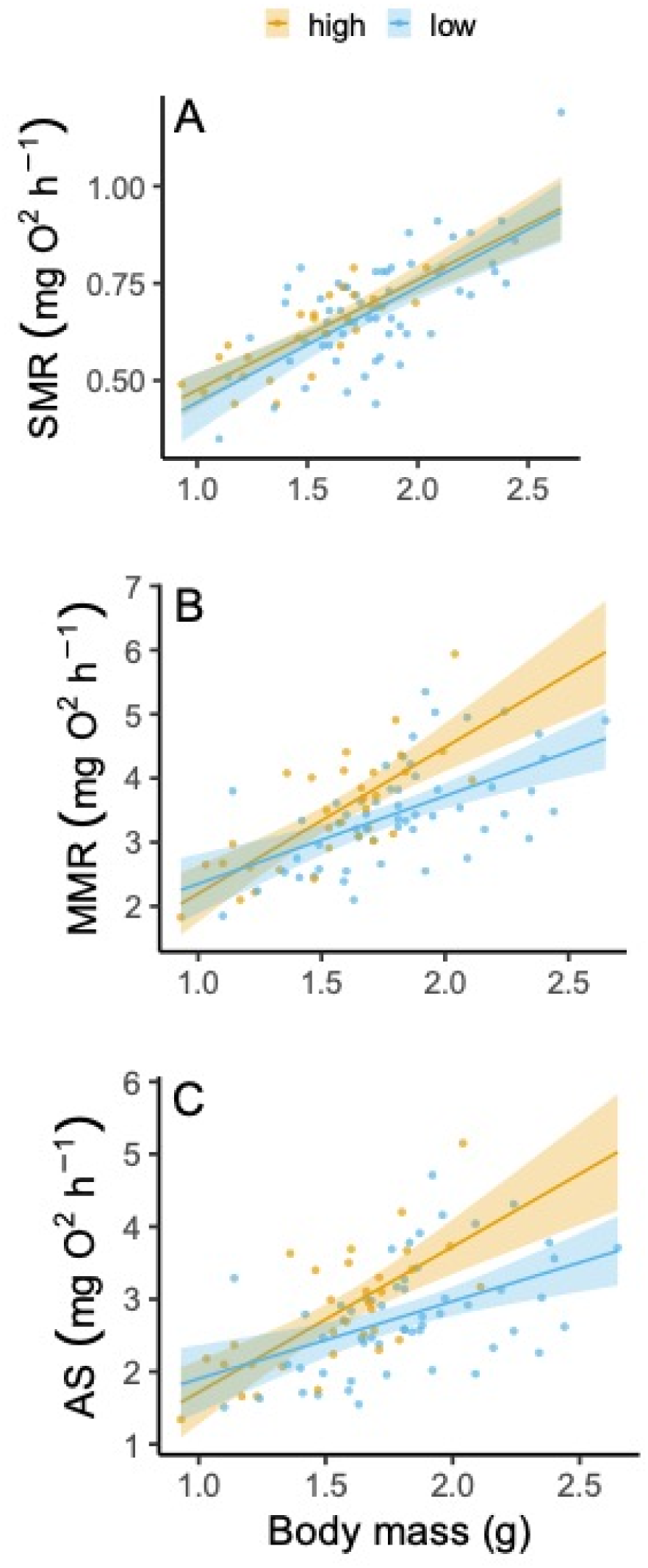
The effect of water flow on the relationship between metabolic rate and body mass within schools of *Chromis viridis*, including (A) standard metabolic rate (SMR), B) maximum metabolic rate (MMR), and (C) aerobic scope (AS). The blue and orange shaded areas illustrate the 95% confidence interval of the regression for low (< 11 cm s^-1^) and high (>20 cm s^-1^) flow sites, respectively, for the relationship between body mass and metabolic rate. Each data point represents an individual fish.

Conversely, SMR did not correlate with water flow rate (χ^2^=1.16, p = 0.281, adjusted R^2^=0.510; Figure 2a), though body mass exerted a significant positive effect (χ^2^=89.96, p < 0.001; Figure 2a). The best-fit model for SMR included only the main effects of water flow rate and mass, without their interaction or a random effect for site.

When comparing between schools within sites, there were no significant differences in any metabolic trait among different schools. For SMR, the best-fit model included both school and body mass as fixed effects, though excluded a school*body mass interaction and a random effect for site. Though the model indicated a significant effect of school on SMR (χ^2^=21.51, p = 0.003, adjusted R^2^=0.644), this effect was driven by differences among schools from different sites, not differences between schools from the same site (Tukey’s multiple comparisons post-hoc test: p > 0.05 for all comparisons of schools within sites). The best-fit models for MMR and AS only included a body mass fixed effect, excluding school, a school*body mass interaction, and a random effect for site. Body mass had a significant effect on metabolic rate for all models, including SMR (χ^2^=92.30, p < 0.001), MMR (χ^2^=38.19, p < 0.001, adjusted R^2^=0.353), and AS (χ^2^=22.41, p < 0.001, adjusted R^2^=0.235).

### Study 2: Mark-recapture of social groups

Out of 280 total tagged fish, 105 individuals were recollected (44%) over the two rounds of collection. Of these 105 fish, 41 had migrated from the school with which they were originally tagged to another school, moving 33m on average (± 3.65 m, s.e.m.; range: 5 – 78 m). Twelve of these fish moved across the sandy patch between Little Vicki’s and No Name reefs, while the remaining 29 migrators remained on the reef in which they were tagged. The body size class of the fish (χ^2^=0.26, p = 0.610, R^2^m=0.045, R^2^c=0.697), recapture frequency (χ^2^=0.67, p = 0.412), and their interaction (χ^2^=0.28, p = 0.595) had no effect on the tendency to move, whose best-fit model was the most complex model.

For fish that did migrate to a new school, the distance and movement type were not related to either body size class, recapture frequency, or their interaction. The best-fit distance moved model included body size class as a fixed effect and school as a random effect (excluding recapture frequency and the body size class*recapture frequency interaction), while the best-fit movement type model was the most complex model. Distance moved was not related to body size class (χ^2^=1.73, p = 0.188, R^2^m=0.044, R^2^c=0.271). Similarly, movement type was not correlated with body size class (χ^2^=1.36, p = 0.243, R^2^m=0.724, R^2^c=0.975), recapture frequency (χ^2^=0.05, p = 0.832), or their interaction (χ^2^=0.00, p = 0.999). However, in nine instances, multiple individuals (two or more) migrated from a shared school of origin to the same destination school (Figure 3).

**Figure 3.**
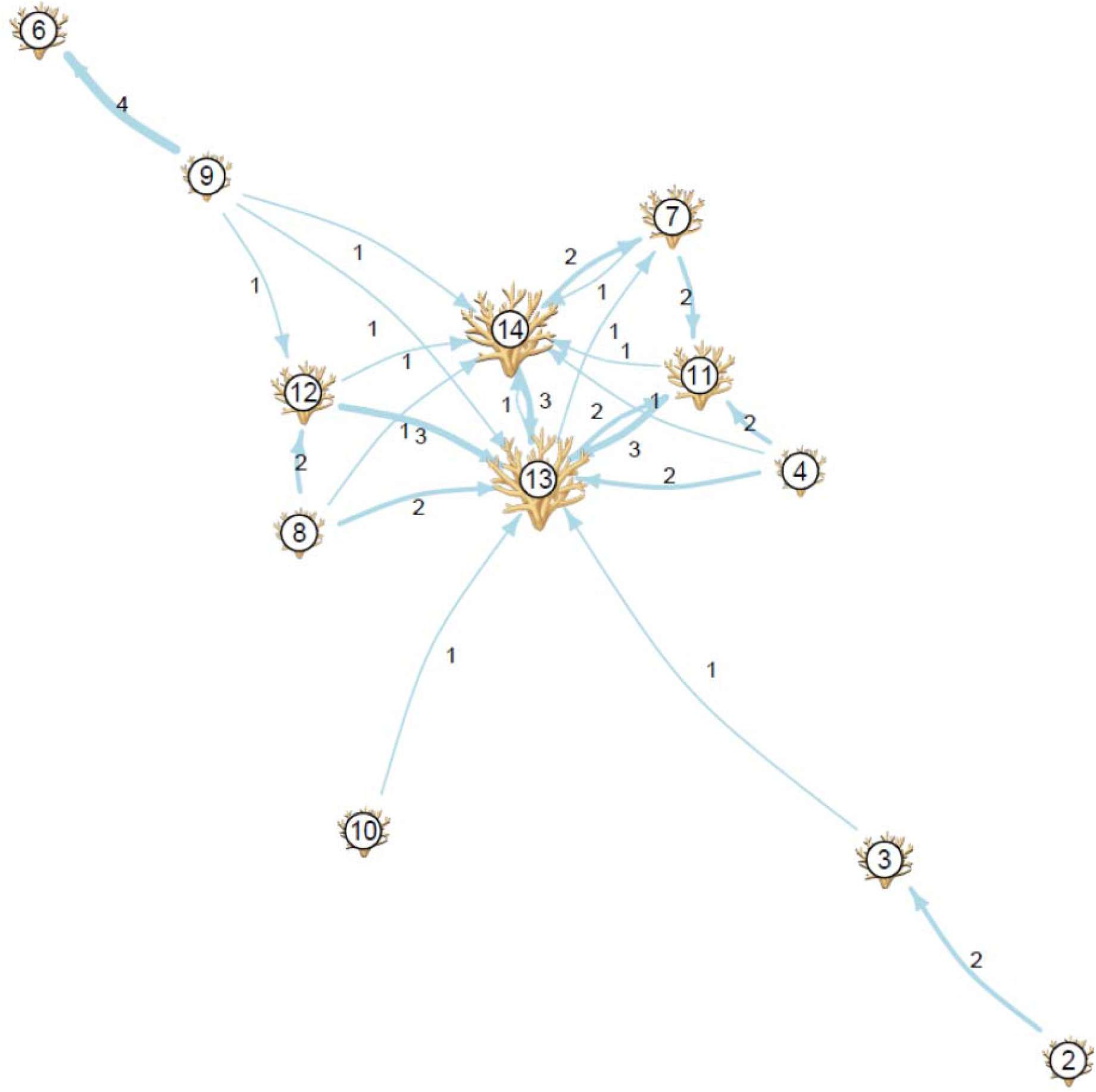
Network diagram showing the connections between schools of the gregarious coral reef fish *Chromis viridis*, visualized using a Fruchterman-Reingold layout. Coral colonies (nodes) were scaled in size according to in-degree centrality, visually emphasizing colonies with a higher influx of fish. Directed edges (blue arrows) indicate the numbers of fish moving between colonies, visualized by arrow thickness and the associated numbered labels beside each arrow. Colonies are numbered to match those shown in the inset of Figure 1.

A school’s social network was modified by both recapture frequency and site (Table S2). Both a school’s degree (i.e., total incoming and outgoing connections) and eigenvector centrality (i.e., which indicated a school’s connectedness in the network) were influenced by site (degree: χ^2^=8.41, p = 0.015, eigenvector centrality: χ^2^=17.57, p < 0.001; Figure 4), as No Name reef had both a higher degree and eigenvector centrality than Little Vicki’s reef, and the school found in the sand between No Name and Little Vicki’s reefs (in which there was only one school found). While neither degree nor eigenvector centrality were influenced by recapture frequency (p > 0.05), eigenvector centrality was impacted by a recapture frequency*site interaction (χ^2^=4.87, p = 0.027; Figure 4b), as schools on No Name reef that were recaptured twice had a higher eigenvector centrality than those that were recollected only once.

**Figure 4.**
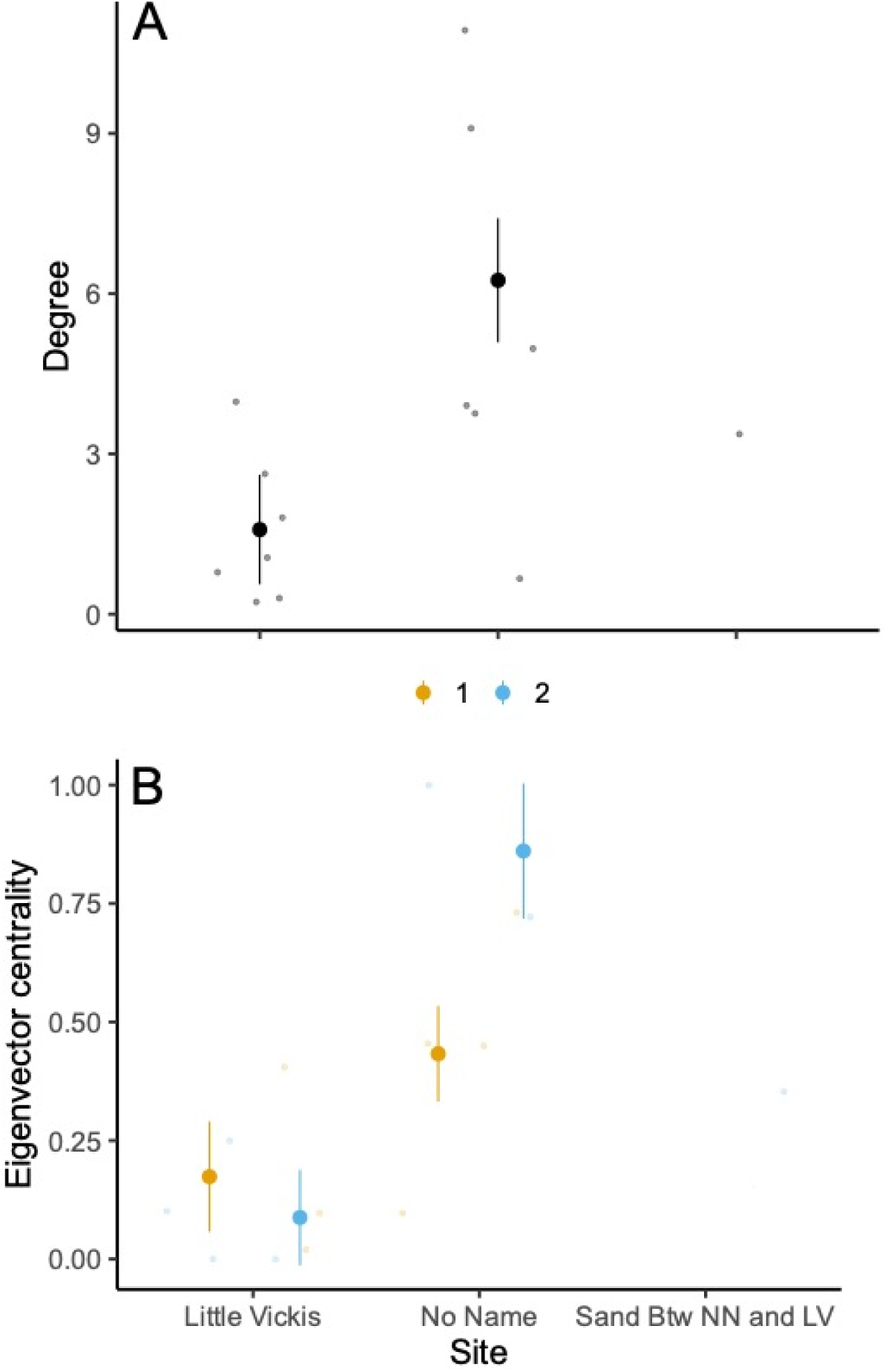
Factors influencing social network metrics, including degree (total incoming and outgoing connections) and eigenvector centrality (a school’s connectedness in the network). (A) Effect of site on node degree (Little Vickis: n=7 schools, No Name: n=6, sand between No Name and Little Vickis: n=1). (B) The interactive effects of site and recapture frequency on eigenvector centrality. Large dots indicate mean ± s.e.m., while small dots indicate individuals.

## Discussion

Our study examined inter-group differences in metabolic traits and variation in the social group fidelity within (i.e., sections of continuous reef) and among (i.e., reefs separated by stretches of sandy substratum) habitats. Importantly, we found that habitats with a higher mean water flow contained groups of damselfish with a higher MMR and AS. However, despite finding that approximately a third of *C. viridis* individuals are migrating among different coral colonies and social groups, no differences were found in any metabolic traits between schools within sites. These findings provide preliminary evidence that, in addition to sex, morphology and behaviour (Rodgers et al. 2011; Cattelan and Griggio 2018; Kelley and Evans 2018; Darden et al. 2020), group composition may also be related to physiological traits associated with energy requirements and locomotor capacity, potentially due to the dynamic and challenging selective pressures imposed by a coral reef environment (Clarke et al. 2009; McCormick et al. 2018).

Several mechanisms could promote heterogeneity in the physiological traits of *C. viridis* social groups among different habitats (Killen et al. 2017b). The first potential mechanism is genetic relatedness. In species in which offspring settle within the same social group as their parents, concordance in phenotype could arise due to similar genetic makeup (Armitage 1987; Charpentier et al. 2010). Although two studies of *Dascyllus* spp. provided evidence of some limited genetic similarity in social groups of coral reef fish with a pelagic larval phase (Buston et al. 2009; Bernardi et al. 2012), most reef species, including *C. viridis*, spend a substantial period of time as a pelagic larvae prior to settling on the reef and entering the juvenile stage (Leis and Carson-Ewart 2000), limiting concurrent settlement of genetic relatives that live within groups of social coral reef fishes (Avise and Shapiro 1986). Instead, connections between physiology and habitat preferences may drive spatial and temporal overlap in settlement of juvenile fishes to the reef through passive assortment mechanisms (Killen 2014), leading to heterogeneity in the distribution of phenotypes in social groups from different habitats (Killen et al. 2017b). Another potential mechanism is phenotypic plasticity, in which individuals within groups change their phenotype so that they can cope with environmental conditions within a site. As *C. viridis* often exhibit high site fidelity to a particular coral colony and social group (with nearly two thirds of fish being recollected from the coral colony on which they were initially tagged in study 2), plasticity throughout development could then enhance differences in phenotypes across broad differences in habitat (e.g. flow regimes), as has been found for other coral reef fishes (Fulton 2010; Fulton et al. 2013; Binning et al. 2014; Binning et al. 2015; Nadler et al. 2018). For example, coral reef fish held in the laboratory under high flow conditions developed an increased capacity for aerobic swimming, including higher MMR and AS (Binning et al. 2015). There could also be a loss of metabolically unsuited individuals at a given site via selective mortality. Traits such as growth rate, size at settlement and post-larval duration influence post-settlement survival in *C. viridis*, but the strength of selection on these traits varies among habitats (Block and Steele 2014). Growth rate and swim performance are often related to AS and MMR in fishes (Claireaux and Lefrancois 2007), and so any selection on these traits could produce correlated selection on metabolic traits in social groups. Given the life-history of *C. viridis,* more than one mechanism is likely responsible for the observed variation in metabolic phenotypes among habitats.

We found no evidence of assortment of metabolic traits within sites, potentially due to several factors. First, there may be no active assortment among social groups within sites, in which individuals select group-mates with a similar physiological phenotype and collectively take up residency in groups that are well suited to their physiological and behavioural traits (Killen et al. 2017b). However, based on the results of study 2, we learned that, despite previous evidence suggesting that *Chromis* spp. exhibit high site fidelity to a single coral colony (Sackley and Kaufman 1996; Chase et al. 2020), *C. viridis* will in fact migrate to new social groups up to 78m away. Thus, our criterion of a minimum of 50m between groups within sites may have been insufficient to ensure that social groups were indeed completely distinct, meaning the individuals may have been engaging in some migration between social groups within sites that prevented detection of within-site assortment. Within a given site, flow regime and bathymetry were also similar, suggesting that phenotypic plasticity and selective mortality were less important drivers of distribution on this small spatial scale (Killen et al. 2017b).

Despite over a third of recollected fish moving to a new social group, movement dynamics of *C. viridis* were not altered by either body size or perceived habitat risk. The trade-offs of migration to a new coral colony or social group are likely dictated by an individual’s environmental limits, habitat needs, or preferences for particular social traits in group-mates. First, for an animal to migrate, it needs to have sufficient locomotor capacity to survive the move, including the ability to overcome any barriers presented by fast or turbulent water flow conditions, with Nadler et al. (2018) showing that schooling fish from higher flow sites have greater locomotor performance than fish from lower flow habitats. Second, at the microhabitat level, coral colony characteristics likely shape habitat preferences for fish with an obligate relationship with the hard coral structure, like *C. viridis*, who are typically found schooling within 30cm of branching coral during the day (presumably feeding) and sleeping within the branches of coral colonies at night (Chase et al. 2020). Given the importance of coral in the life history strategy for coral-dwelling fishes, it is unsurprising that these species show clear preferences by both coral species and coral colony traits that may promote migration (Kane et al. 2008; Schrandt et al. 2012; Nadler et al. 2018; Chase and Hoogenboom 2020). Third, the social composition of a group alters its suitability for individuals. Killen et al. (2021a) illustrated the connection between an individual’s metabolic rate and its sociability in the coral reef fish species, the red belly yellow tail fusilier *Caesio cuning*, finding that individuals with a higher MMR were more sociable than conspecifics with a lower MMR. Migration may also be more favourable if an individual does not need to embark on a potentially risky journey alone. In the humbug damselfish *Dascyllus aruanus*, the tendency to move between habitats is dictated by the behaviour of multiple group members, requiring both an initiating leader to start the move and willingness of group members to follow (Ward et al. 2013). In this study, we cannot confirm that migrations occurred in groups, but on multiple occasions, we found two or more fish from the same original school with the same new school, suggesting that these migrations may, on several occasions, have been completed in pairs or groups. Testing the nature of migrations (i.e., whether they were completed in a solitary or social context) would form an ideal follow up study to this one, to better understand the importance of social interactions to the trade-offs of social group migration.

Recently, research has focused on the role of intraspecific variation in behavioural and physiological traits and understanding the factors that generate and maintain this variation (reviewed in Cope et al. 2022; Winberg and Sneddon 2022). Our results suggest that phenotypic differences among social groups in the wild may be an important consideration when quantifying intraspecific variation and the distribution of phenotypes across habitats. The general picture emerging from the current study is that, in a damselfish species with high affinity for live coral structure and frequently high site fidelity (as nearly 2/3 of recollected fish were found on the same coral colony), a combination of mechanisms likely dictates the observed metabolic phenotypes in social groups and tendency to migrate among coral colonies and social groups. Assortment, selective mortality, and phenotypic plasticity all potentially contributing in varying degrees at different spatial scales (Killen et al. 2017b). The physiological performance of group mates will play an important role in modulating outcomes during events which have the greatest impact on individual fitness, such as during predator attacks or exposure to stressful environments (Nadler et al. 2018; Tiddy et al. 2024). While these data provide the first evidence to support the distribution of social groups according to physiological characteristics, additional work is needed to understand how physiology shapes association preferences among social fishes, the mechanisms responsible for physiological assortment, and its importance for structuring intra-specific phenotypic diversity on a variety of spatial scales.

## Acknowledgements

Thank you to the staff at the Lizard Island Research Station, Katherine Corkill, Rahel Zemoi, and Stephen Brown for logistical support. This research was conducted under James Cook University Animal Ethics approval number A2103. Research funding was provided by a Natural Environment Research Council Advanced Fellowship (SSK, NE/J019100/1), Australian Postgraduate Award, International Postgraduate Research Scholarship, Lizard Island Reef Research Foundation Doctoral Fellowship, Great Barrier Reef Marine Park Authority Science for Management Award and James Cook University Graduate Research Scheme (LEN). All data and code are available in the University of Southampton Institutional Research Repository (DOI available once paper is accepted for publication).

## Notes

### Competing Interest Statement

The authors have declared no competing interest.

