## Supplementary tables for "Evidence of physiological assortment and movement dynamics among social groups of a coral reef fish"

**Submitted to *Coral Reefs***

[Lauren E. Nadler^1,2*†^](https://orcid.org/0000-0001-8225-8344), [Mark I. McCormick^3^](https://orcid.org/0000-0001-9289-1645), Amy Cox^4^, [Kathryn Grazioso^5,6^](https://orcid.org/0009-0009-5140-2396), [Shaun S. Killen^7†^](https://orcid.org/0000-0003-4949-3988)

^1^ College of Marine and Environmental Sciences, ARC Centre of Excellence for Coral Reef Studies, James Cook University, Townsville, QLD, 4811, Australia

^2^ School of Ocean and Earth Science, National Oceanography Centre, University of Southampton, Southampton, SO14 3ZH, United Kingdom

^3^ Coastal Marine Field Station, School of Science, University of Waikato, Tauranga 3110, New Zealand

^4^ Biological and Chemical Sciences, Cave Hill Campus, The University of the West Indies, Bridgetown BB11000, Barbados

^5^ NOAA, Southeast Fisheries Science Center, National Marine Fisheries Service, Miami, FL, 33149, USA

^6^ University of Miami, Cooperative Institute of Marine & Atmospheric Studies, Miami, FL, 33149, USA

^7^ Institute of Biodiversity, Animal Health & Comparative Medicine, College of Medical, Veterinary & Life Sciences, University of Glasgow G12 8QQ, United Kingdom

†These authors contributed equally to this manuscript

**TABLE S1.** **Checklist of 53 essential criteria for the reporting of methods for aquatic intermittent-flow respirometry**. (This table can be copied by authors and included as supplemental information when submitting manuscripts and publishing papers. This will facilitate clear and concise means for reporting all important methodological details, with the article main text being used to provide additional information and context.)

| Number | **Criterion and Category** | **Response** | **Value (where required)** | **Units** |
| --- | --- | --- | --- | --- |
|  | **EQUIPMENT, MATERIALS, AND SETUP** |  |  |  |
| 1 | Body mass of animals at time of respirometry | mean | 1.717 | g |
| 2 | Volume of empty respirometers |  | 70 | ml |
| 3 | How chamber mixing was achieved | peristaltic pump |  |  |
| 4 | Ratio of net respirometer volume (plus any associated tubing in mixing circuit) to animal body mass | 44:1 |  |  |
| 5 | Material of tubing used in any mixing circuit | Tygon gas impermeable |  |  |
| 6 | Volume of tubing in any mixing circuit |  | 5 | ml |
| 7 | Confirm volume of tubing in any mixing circuit was included in calculations of oxygen uptake | Yes |  |  |
| 8 | Material of respirometer (e.g. glass, acrylic, etc.) | glass with acrylic end caps |  |  |
| 9 | Type of oxygen probe and data recording | Pyroscience optodes with Firesting unit |  |  |
| 10 | Sampling frequency of water dissolved oxygen | Every 2 seconds |  |  |
| 11 | Describe placement of oxygen probe (in mixing circuit or directly in chamber) | In mixing circuit |  |  |
| 12 | Flow rate during flushing and recirculation, or confirm that chamber returned to normoxia during flushing | Chamber returned to normoxia during flushing |  |  |
| 13 | Timing of flush/closed cycles | 2 min flush/7 min closed |  |  |
| 14 | Wait (delay) time excluded from closed measurement cycles |  | 30 | seconds |
| 15 | Frequency and method of probe calibration (for both 0 and 100% calibrations) | Both at beginning of study, daily for 100% |  |  |
| 16 | State whether software temperature compensation was used during recording of water oxygen concentration | No temperature compensation |  |  |
|  | **MEASUREMENT CONDITIONS** |  |  |  |
| 17 | Temperature during respirometry |  | 28.74 | ^o^C |
| 18 | How temperature was controlled | Heated reservoir |  |  |
| 19 | Photoperiod during respirometry | Approximately 12h/12h |  |  |
| 20 | If (and how) ambient water bath was cleaned and aerated during measurement of oxygen uptake (e.g. filtration, periodic or continuous water changes) | Water flowing from reservoir through the water bath continuously, aerated with air stone. |  |  |
| 21 | Total volume of ambient water bath and any associated reservoirs | Unknown |  |  |
| 22 | Minimum water oxygen dissolved oxygen reached during closed phases |  | 80% | Air saturation |
| 23 | State whether chambers were visually shielded from external disturbance | Yes |  |  |
| 24 | How many animals were measured during a given respirometry trial (i.e. how many animals were in the same water bath) |  | 8 |  |
| 25 | If multiple animals were measured simultaneously, state whether they were able to see each other during measurements | No |  |  |
| 26 | Duration of animal fasting before placement in respirometer |  | 24 | hours |
| 27 | Duration of all trials combined (number of days to measure all animals in the study) |  | 12 | days |
| 28 | Acclimation time to the laboratory (or time since capture for field studies) before respirometry measurements |  | 3-5 | days |
|  | **BACKGROUND RESPIRATION** |  |  |  |
| 29 | State whether background microbial respiration was measured and accounted for, and if so, method used (e.g. parallel measures with empty respirometry chamber, measurements before and after for all chambers while empty, both) | Oxygen uptake in all chambers was measured while empty both before and after each trial, and all measures of oxygen uptake were corrected assuming a linear increase in bacterial oxygen consumption with time. |  |  |
| 30 | State if background respiration was measured at beginning and/or end, state how many slopes and for what duration | Measured at beginning at end, three slopes of 7 min each. |  |  |
| 31 | State how changes in background respiration were modelled over time (e.g. linear, exponential, parallel measures) | Linear. |  |  |
| 32 | Level of background respiration (e.g. as a percentage of SMR) |  | 17.03 | % |
| 33 | Method and frequency of system cleaning (e.g. system bleached between each trial, UV lamp) | Every morning the system was cleaned with soap and hot water. |  |  |
|  | **STANDARD OR ROUTINE METABOLIC RATE** |  |  |  |
| 34 | Acclimation time after transfer to chamber, or alternatively, time to reach beginning of metabolic rate measurements after introduction to chamber |  | 2 | hours |
| 35 | Time period, within a trial, over which oxygen uptake was measured (e.g. number of hours) |  | ~18 | hours |
| 36 | Value taken as SMR/RMR (e.g. quantile, mean of lowest 10 percent, mean of all values) |  | 10th | percentile |
| 37 | Total number of slopes measured and used to derive metabolic rate (e.g. how much data were used to calculate quantiles) |  | ~130 |  |
| 38 | Whether any time periods were removed from calculations of SMR/RMR (e.g. data during acclimation, periods of high activity [e.g. daytime]) | First two hours after transfer to chambers |  |  |
| 39 | r^2^ threshold for slopes used for SMR/RMR (or mean) |  | 0.95 |  |
| 40 | Proportion of data removed due to being outliers below r-squared threshold |  | <5% |  |
|  | **MAXIMUM METABOLIC RATE** |  |  |  |
| 41 | When MMR was measured in relation to SMR (i.e. before or after) | Before |  |  |
| 42 | Method used (e.g. critical swimming speed respirometry, swim to exhaustion in swim tunnel, or chase to exhaustion) | Chase to exhaustion |  |  |
| 43 | Value taken as MMR (e.g. the highest rate of oxygen uptake value after transfer, average of highest values) | Highest rate after transfer, taken during first slope of at least 10 minutes, during any 3 minute window during this time. |  |  |
| 44 | If MMR measured post-exhaustion, length of activity challenge or chase (e.g. 2 min, until exhaustion, etc.) |  | 90.07 average | seconds |
| 45 | If MMR measured post-exhaustion, state whether further air-exposure was added after exercise | Yes | 30 | seconds |
| 46 | If MMR measured post-exhaustion, time until transfer to chamber after exhaustion or time to start of oxygen uptake recording |  | <10 | seconds |
| 47 | Duration of slopes used to calculate MMR (e.g. 1 min, 5 min, etc.) |  | 3 | minutes |
| 48 | Slope estimation method for MMR (e.g. rolling regression, sequential discrete time frames) | sequential discrete |  |  |
| 49 | How absolute aerobic scope and/or factorial aerobic scope is calculated (i.e. using raw SMR and MMR, allometrically mass-adjusted SMR and MMR, or allometrically mass-adjusting aerobic scope itself) | Difference between raw MMR and SMR. |  |  |
|  | **DATA HANDLING AND STATISTICS** |  |  |  |
| 50 | Sample size | N = 8 fish from each of 11 shoals (88 fish in total) |  |  |
| 51 | How oxygen uptake rates were calculated (software or script, equation, units, etc.) | Fishresp R Script |  |  |
| 52 | Confirm that volume (mass) of animal was subtracted from respirometer volume when calculating oxygen uptake rates | Yes |  |  |
| 53 | State whether analyses accounted for variation in body mass and describe any allometric mass-corrections or adjustments | Body mass included as a covariate in models. |  |  |

**Table S2.** Summary of social network metrics. To quantify the connectivity and movement dynamics of fish in schools on coral colonies (‘colony’), we calculated the following network metrics for each colony: 1) in-degree: representing the total number of fish arriving at a colony from other colonies; 2) out-degree: representing the total number of fish leaving a colony to other colonies; 3) total degree: the sum of in-degree and out-degree; 4) betweenness: quantifies the role of a colony as an intermediary or "bridge" in the network; and 5) eigenvector centrality: summarises direct connections of a colony and the centrality of its neighbours.

| Colony | Total degree | In_degree | Out-degree | Betweenness | Eigenvector centrality |
| --- | --- | --- | --- | --- | --- |
| 2 | 1 | 0 | 1 | 0 | 0.01955656 |
| 3 | 2 | 1 | 1 | 4 | 0.10080048 |
| 4 | 3 | 0 | 3 | 0 | 0.40492351 |
| 6 | 1 | 1 | 0 | 0 | 0.09683984 |
| 7 | 4 | 2 | 2 | 1.5 | 0.44969192 |
| 8 | 3 | 0 | 3 | 0 | 0.3531023 |
| 9 | 4 | 0 | 4 | 0 | 0.2495711 |
| 10 | 1 | 0 | 1 | 0 | 0.09700626 |
| 11 | 5 | 3 | 2 | 0 | 0.72172476 |
| 12 | 4 | 2 | 2 | 0 | 0.45462236 |
| 13 | 11 | 8 | 3 | 13.5 | 1 |
| 14 | 9 | 7 | 2 | 4.5 | 0.73075011 |
